# Mesenchymal Stromal Cell Delivery via an Ex Vivo Bioreactor Preclinical Test System Attenuates Clot Formation for Intravascular Application

**DOI:** 10.1101/2020.11.20.391631

**Authors:** Brian O’Rourke, Sunny Nguyen, Arno W. Tilles, James A. Bynum, Andrew P Cap, Biju Parekkadan, Rita N. Barcia

## Abstract

While mesenchymal stromal cells (MSCs) are an appealing therapeutic option for a range of clinical applications, their potential to induce clotting when used systemically remains a safety concern, particularly in hypercoagulable conditions, such as in patients with severe COVID-19, trauma, or cancers. Here, we tested a novel *ex vivo* approach aimed at improving the safety of MSC systemic administration by use of a bioreactor. In this device, MSCs are seeded on the outside of a hollow-fiber filter, sequestering them behind a hemocompatible membrane, while still maintaining cross talk with blood cells and circulating signaling molecules. The potential for these bioreactor MSCs to induce clots in coagulable plasma was compared against “free” MSCs, as a model of systemic administration, which were directly injected into the circuit. Our results showed that physical isolation of the MSCs via a bioreactor extends the time necessary for clot formation to occur when compared to “free” MSCs. Measurement of cell surface data indicates the presence of known clot inducing factors, namely tissue factor and phosphatidylserine. Results also showed that recovering cells and flushing the bioreactor prior to use further prolonged clot formation time. Further, application of this technology in two *in vivo* models did not require additional heparin to maintain target ACT levels relative to the acellular device. Taken together, the use of hollow fiber filters to house MSCs, if adopted clinically, could offer a novel method to control systemic MSC exposure and prolong clot formation time.

## Introduction

Mesenchymal stromal cells (MSCs) are potent immunoregulators with strong preclinical data that support their application in a wide range of clinical conditions [1–4]. MSCs can provide therapeutic benefit to patients suffering from systemic inflammation by effectively immunomodulating peripheral blood cells to reduce inflammatory signaling and promote homeostasis. Significant evidence of these MSC derived effects have been shown *in vitro*, in animal models, and in clinical trials, including recently under an emergency IND application or expanded access protocol with COVID-19 infected patients [5–10]. However, it is known that under certain conditions MSCs may be pro-coagulable and promote instant blood-mediated inflammatory reaction (IBMIR), likely through the expression of known coagulation factors on their cell surfaces and in the production of cellular microvesicles [11–14]. Two such factors, tissue factor and phosphatidylserine are known to be integral in physiological coagulation [15–18]. Phosphatidylserine is a major component of cell-based coagulation, enhancing coagulation activity through the charge-based binding of coagulation factor zymogens and cofactors to increase formation of the tenase and prothrombinase complexes [19–21]. This binding effectively enhances clot formation potential, exacerbating a response when a physiological trigger of coagulation, such as tissue factor, is present [18, 21, 22]. When MSCs are introduced into systemic circulation they bring with them the tissue factor expressed on their cell surface. MSCs sourced from different donors and locations (bone marrow, BM-MSCs, adipose-derived, AD-MSCs, umbilical cord, UC-MSCs) vary in tissue factor expression, with higher levels of tissue factor correlating with quicker clot formation and IBMR. BM-MSCs, the most commonly used, have been widely shown to have the least tissue factor expression [16, 18].

Reducing potential MSC driven clot formation has been a major focus for both academic and clinical groups over the last decade. Advances have been made to mitigate these risks including changing the MSC mode of delivery away from systemic exposure via intravenous (IV) infusion towards localized injection. However, in some cases, systemic IV infusions could provide the highest therapeutic potential. Minimizing clot formation potential could improve therapeutic efficacy, and also increase safety of the treatment in hypercoagulable patients such as patients with COVID-19 in the ICU [23–26]. To that end, increasingly rigorous release criteria during MSC clinical manufacturing are now being applied. Cell populations with low tissue factor are being selected and ‘fresh’ or recovered MSCs with lower surface phosphatidylserine exposure are preferred [17, 27]. Additional clot mitigation approaches are being evaluated, including shifting delivery of the MSCs from intravenous to intramuscular administration, emphasizing higher cell viability, or even using gene modification to promote cell survival [28–31].

We propose an alternative approach which would minimize direct exposure of the MSCs to blood and contain MSCs in one location. We utilized a recently described perfusion platform that incorporates a hollow-fiber filter into a fluid circuit to compartmentalize the MSCs, while still allowing exchange of signaling molecules from perfusate to cells, and vice-versa [32]. MSCs within this platform were shown to effectively retain their immunomodulatory capacity and alter perfused lymphocyte proliferation, activation, and cytokine production in an MSC dose and duration exposure dependent manner, despite having minimal direct contact with the blood cells.

Benchtop coagulation assays, including microfluidic setups, are becoming increasingly translationally relevant[33]. Here, we used a MSC bioreactor platform to assay whether limiting the direct exposure of fresh frozen pooled plasma from healthy patients to MSCs, and therefore the available tissue factor and phosphatidylserine of the MSCs, would affect clot formation time (CFT) in a modified plasma-based clot formation assay [34, 35]. Plasma was perfused through bioreactors seeded with MSCs as well as through circuits in which cells were directly injected into the perfused plasma to make comparative CFT measurements. We were able to show that bioreactor use significantly prolonged CFT relative to direct injection of the MSCs. Flushing of concentrated soluble factors from bioreactors further contributed to prolonged CFTs. Lastly, the previously proposed clinical solution for MSC driven clot formation, anticoagulation with heparin [36–39], was shown to be effective in both perfusion setups. These results suggest this new modality for systemic MSC delivery may offer a safer alternative to intravenous MSC injection. The clinically scaled ex vivo engineered MSC delivery is currently undergoing testing in a Phase I/II trial in acute kidney injury (AKI) and COVID-19 associated AKI.

## Materials and Methods

### MSC Cell Source and Culture Processes

Human bone marrow derived mesenchymal stromal cells were isolated from 3 separate donors. Cells were cultured and propagated either in-house using proprietary techniques at developed Sentien Biotechnologies (MA, USA) or RoosterBio (MD, USA) and cryopreserved at early passage (P3-P5). Cells were cultured using 2D planar growth conditions. Cells from Sentien Biotechnologies were cultured in aMEM media (HyClone, UT, USA, SH3A5195) containing FGF and FBS, while Roosterbio cells were expanded in xeno-free RoosterNourish-MSC-CC media (RoosterBio, MD, USA, KT-021). All human samples were obtained from commercial vendors under a consented protocol for research purposes only.

### MSC Direct Injection

MSCs were thawed from cryopreservation into citrated fresh-frozen pooled plasma (FFPP) (George King Bio-Medical, KS, USA) and counted via Trypan Blue exclusion. Each direct injection subgroup was processed individually and subgroups were never combined. The fresh thawed direct injection subgroup was placed into plasma after counting and directly injected for perfusion (Figure 2). The recovered direct injection subgroup allowed for 24 hours of recovery on a cell culture dish, followed by dissociation with TryPLE, counting via Trypan Blue exclusion, and placing into plasma prior to direct injection. Cells for use in washed groups were washed with saline directly after thaw, pelleted and resuspended in fresh citrated media prior to direct injection.

### Bioreactor Fill/Finish

MSCs were thawed from cryopreservation into aMEM supplemented with 10% HSA and counted via Trypan Blue exclusion prior to seeding in device. The desired cell number was suspended into 9 mL of aMEM and then seeded into saline-primed microreactors (Spectrum Laboratories, CA, USA; C02-P20U-05) with 0, 1, or 3 ×10^6^ viable cells per device (0M, 1M, 3M, respectively). Excess media flowed through the semi-permeable hollow fibers while cells remained within the extraluminal space of the reactor. Microreactors used were 20 cm long with an internal surface area of 28 cm^2^. Within each microreactor are nine 0.5 mm diameter fibers comprised of polyethersulfone with a 0.2 μm pore size. The total internal volume of the microreactor is 1.5 mL.

Depending on the group, microreactors were either used immediately or incubated at 37°C for 2 hours to allow for cell attachment and were subsequently held for24 hours at room temperature prior to integration into the circuit. This hold time intends to mimic the time between device manufacture and potential clinical application.

Following hold, select MRs were subjected to flushing. Prior to connection to the perfusion circuit sterile saline (4.5 mL) was pushed via syringe through the extracapillary port of the MR. Discharge exited through the intracapillary port.. Samples from the extracapillary space were collected prior to and post flush to measure soluble levels of phosphatidylserine and tissue factor.

Large scale bioreactors were used in the animal studies. Similar to the microreactor fill/finish process, following MSC thaw, cell and media suspension was perfused through the extracapillary port onto semi-permeable hollow fibers (Asahi Kasei Medical Inc, IL, USA, OP-05W(A). Cells remained within the extraluminal space of the reactor while excess media perfused through the membranes out the intracapillary port. Thawed vials used in these assays were comprised of cells with a minimum 80% viability. Bioreactors were seeded with either 0, 250, or 750 ×10^6^ viable cells per unit.

### Fresh-Frozen Pooled Plasma

Fresh-Frozen Pooled Plasma was collected and citrated via FDA licensed blood centers from prescreened healthy donors (Geroge King Bio-Medical, KS, USA). No buffers or stabilizers were added. Plasma was frozen within 30 minutes of collection at −70°C from a pool of >50 donors per lot. This plasma still contains many essential factors for clot initiation, including prothrombin which can be activated with the addition of calcium via CaCl_2_ to form firm clots over time. Testing was done to ensure normal values for PT, aPTT, fibrinogen, dRVVT normalized ratio, Factors II, V, VII, VIII, IX, X, XI, and XII.

### Plasma Perfusion

After thawing 5 mL of FFPP via water bath (37°C), 50 uL of 1M CaCl_2_ was added to the plasma within a capped syringe and inverted. The plasma was loaded into prepared perfusion circuits with (0, 1, and 3×10^6^ MSCs) and without microreactors via the syringe port and perfused at a flow rate of 1 mL/min for 5 minutes. Plasma was then extracted from the circuit via the syringe port, aliquoted into microplate wells and placed within the spectrophotometer (Synergy Mx, BioTek) for reading at 405 nm every 10 seconds for a total of 45 minutes.

Positive controls of Innovin (Siemens Healthcare Diagnostics, Germany) or Factor IXa (Haematologic Technologies, VT, USA) were used separately, where designated, at 1:50,000. These positive controls were added prior to CaCl_2_ addition to ensure equal mixing before coagulation initiation. 1M CaCl_2_ was added at 1:100.

Unfractionated heparin (Grifols, Spain) was used at 1.5 U/mL in the designated groups. Heparin was added prior to CaCl_2_ addition to ensure equal mixing before coagulation initiation.

### Clot Formation Time Analysis and Graphing

Clot formation time measurements incorporated the sum of two time periods. The first period initiates when plasma has been recalcified through the addition of CaCl_2_ and continues through perfusion until transfer of the samples into microwells for analysis on the spectrophotometer. This value is immediately recorded by the operator. The second time period occurs when the spectrophotometric readings begin and continues for 45 minutes. At the completion of the study the elapsed time in the first period is added to the time required to obtain the ½ maximal spectrophotometric value, as determined using the clot formation time formula. Combined, these measurements capture the clot formation time.

Resulting values were then graphed and statistically analyzed using unpair student’s t-test (GraphPad Software, La Jolla, CA). Results are presented as mean ± standard deviation. Values of p < 0.05 were considered statistically significant for all analyses.

### Flow cytometry

To measure tissue factor by flow cytometry, staining was done using CD142-APC monoclonal antibody (eBiosciences) in a total volume of 100 μL Stain Buffer containing FBS and ≤0.09% sodium azide (BD Biosciences). Samples were incubated for 15 min at 4°C and analyzed on a FACSCanto II flow cytometer (BD Biosciences) using BD FACSDiva v6.1.1 software. Mouse PE IgG1 kappa isotype antibody (eBioscience) was used as a negative control.

Annexin V staining was done using the FITC Annexin V Apoptosis Detection Kit (BioLegend) according to the manufacturer’s instructions. Fresh thawed MSCs were used to optimize fluorescence compensation.

Flow cytometry analysis was performed in FlowJo (FlowJo LCC, OR, USA; version 10.7).

### Mongrel Dog Perfusion

Eighteen male mongrel dogs were randomized onto the study and underwent surgery for placement of a dialysis catheter (Toxikon Corp, Beford MA). Animals received buprenorphine (0.01 mg/kg, IM) pre-surgery, PM the day of surgery and AM the day after surgery, and cefazolin (22 mg/kg, IV/IM) pre-surgery, then daily for two additional days post-surgery. After a 2 day wash out period, animals were dosed according to their group assignment with 6 animals assigned to either a 0M, 250M, or 750M MSC dose group. Heparin was administered throughout treatment with all animals first receiving a bolus of heparin at 150 U/kg and then a continuous infusion of 25 U/kg every hour. All animals underwent a 24-hour perfusion (+/- 1 hour), with the exception of animal 1003 (Group 1, Control) which was stopped after 22 hours due to low blood flow rate through the catheter.

### Porcine Acute Myocardial Infarction Model Perfusion

On Day 0, 8 Yorkshire pigs underwent induced myocardial infarction of the anterior/septal left ventricle by 45-minute occlusion of the left anterior descending artery (CBSet Inc., Lexington, MA). After one hour of reperfusion/stabilization, animals were connected to the extracorporeal loop via the jugular vein which enabled whole blood circulation through a large scale bioreactor for a period of up to 12 hours. Bioreactors were seeded with either 0 or 750 million human bone-marrow derived MSCs, 24 hours prior to perfusion (n=4 per group). Heparin was administered throughout the treatment, first as a bolus of 225 U/kg and then intermittently to ensure ACTs remained above 300 seconds. Serum troponin levels were assessed at baseline 12 and 24 hours after infarction induction. At 72 hours post injury induction animals were sacrificed with their hearts excised and dyed with Evans Blue and 1% TCC. Infarct area was determined through tracing of digitized images of section via the morphometric software system Olympus cellSens (Version 1.17).

## Results

### Development of an assay to test CFT under perfusion

We developed an approach in which we could test human plasma compatibility of allogeneic MSCs across multiple extents of cell exposure. Cryopreserved MSCs could either be injected into the plasma directly after thaw (**Fig. 1A**), cultured for 24 hours after recovery and then directly injected into plasma (**Fig. 1B**), or seeded into hollow-fiber microreactors with a semi-permeable membrane, allowed to attach, and held for up to 24 hours before being subjected to perfusion (**Fig. 1C**). These ranges of administration broadly represent many of the systemic administration options available today and allow the comparison of varying degrees of MSC to plasma exposure and the effects of MSC culture conditions.

**Figure 1.**
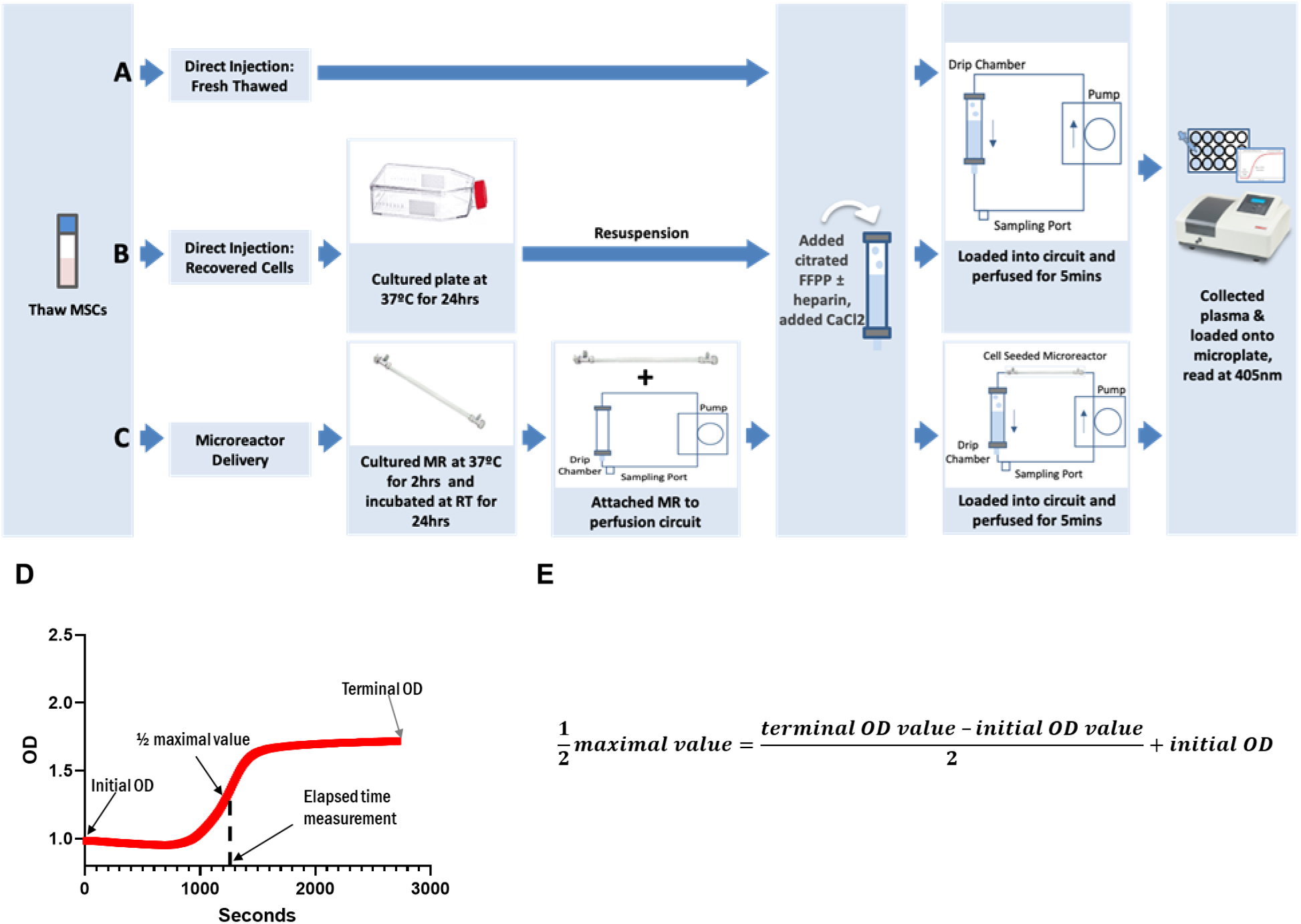
Clot Formation Assay Experimental Setup. MSCs were thawed from cryopreservation and were either **(A)** immediately combined into fresh-frozen pooled plasma-based (FFPP), perfused in the circuit and read, **(B)** cultured for 24 hours then resuspended into FFPP, perfused in the circuit and read, or **(C)** seeded into micro bioreactors (MR), allowed to attach to the hollow-fiber filters for 2 hours, room temperature incubated for 24 hours then attached to perfusion circuits loaded with FFPP, perfused and read. Perfusion of the MR circuits lasted 5 minutes before samples were collected and read on the spectrophotometer at 405nm to assess fibrin formation. (**D**) Resulting spectrophotometric optical density (OD) readouts were graphed over time. (**E**) Formula used for the calculation of the clot formation time determined at the ½ maximal value.

After perfusion, plasma was collected from the circuit via the syringe port and placed into microwells for spectrophotometric reading over time. The point at which a clot was formed was determined by using the resultant spectrophotometric readout and formula (**Fig. 1D, E**) to calculate the ½ maximal value, a point previously determined to designate clot formation [34, 35]. Higher values for clot formation times indicate slower clot formation within the plasma.

### Presence of MSCs accelerates CFT

Consistent with previous work [12], our fresh-frozen pooled plasma-based (FFPP) clot formation assay showed that the presence of MSCs accelerated clot formation under flow conditions relative to acellular controls (**Figure 2**). Interestingly, the same experimental setup run with plasma isolated from an individual donor 24 hours after collection instead of FFPP did not result in significantly different clotting times between cellular and acellular groups, likely as it wasn’t frozen directly after being pulled (**Figure S1**). Spectrophotometric measurements of optical density at 405 nm captured fibrin clot formation as it occurred within the microwell via increases in measured OD over time (**Figure 2A**). Results from our assays showed that the direct injection (DI) of freshly thawed MSCs into the plasma flow circuit significantly hastened the onset of clot formation when compared to circuits using bioreactor housed MSCs. Both the 1M BM-MSC and 3M BM-MSC direct injection groups induced clot formation during the initial perfusion, prior to spectrophotometric reading. All other groups completed perfusion and spectrophotometric reading (**Figure 2B**). These findings were consistent across 3 separate MSC donors (**Figure S2**).

**Figure 2.**
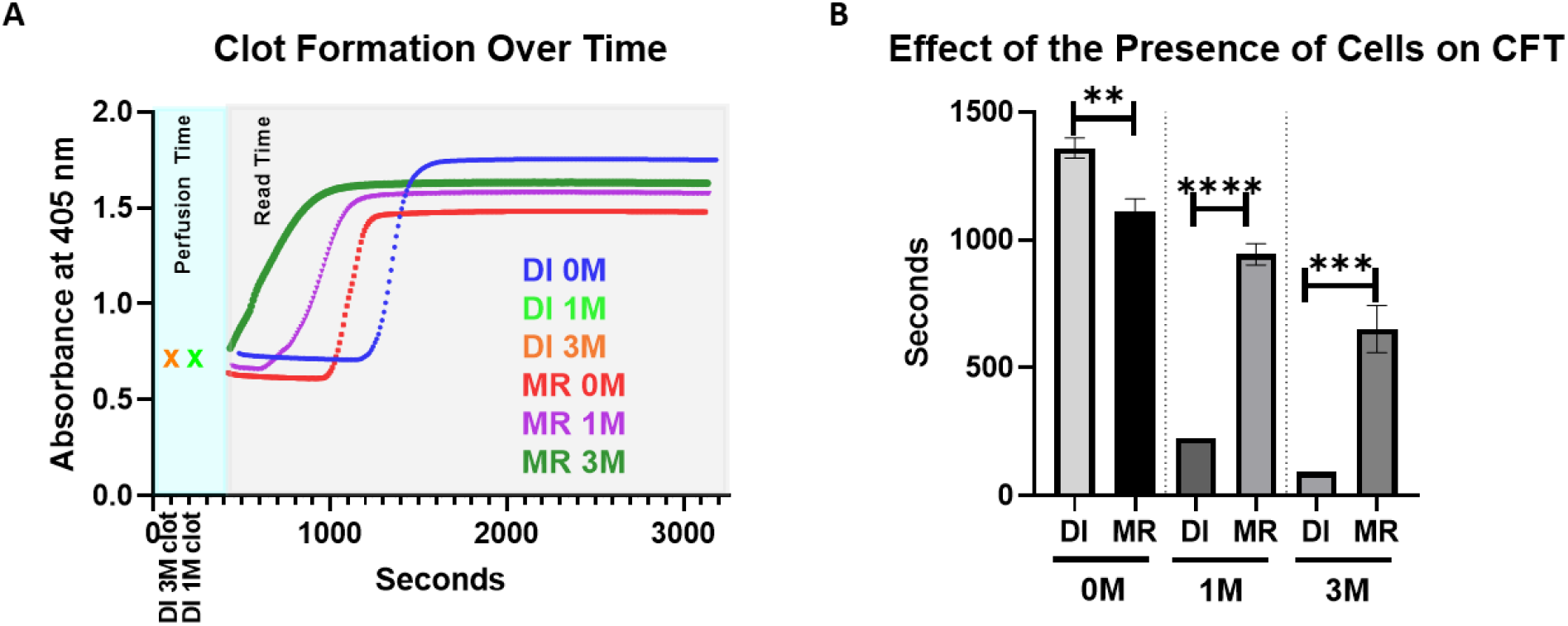
Increased MSC Exposure Shortens Clot Formation Time. 0, 1 x 10^6^, or 3 x 10^6^ viable MSCs were isolated for inclusion in each respective group. Freshly thawed MSCs were used in direct injection (DI) groups. MSCs used in MR groups were first incubated for 2 hours at 37 °C followed by a 24 hour hold at room temperature prior to perfusion. After each groups’ cells were prepared warmed fresh frozen pooled plasma was perfused through circuit for 5 minutes then subjected to spectrophotometric measurements. (**A**) Measurements of fibrin clot formation in plasma were made every 10 seconds over a 45-minute period (grey shaded region) following 5 minutes of perfusion (aqua shaded region). Groups which clotted during perfusion are designated with an ‘x’ at the time at which the clot was noted to be visibly obstructing perfusion. As clots formed, absorbance increased resulting in the designated curves. (**B**) Values for CFT were determined. Resulting values were graphed and analyzed with an unpaired student’s t-test. N=3 runs per group. **=p<0.005; ***= p= 0.0005; ****= p< 0.0001. Error bars represent ± standard deviation. DI = direct injection

Furthermore, MSC dose played a role in clot formation. Increases in the number of cells administered were significantly associated with shorter CFTs across both administration routes. However, when comparing similarly seeded MRs to direct injection with the same number of cells, CFT was significantly slower in the groups where MSCs were housed in the bioreactor (**Figure 2B**).

### Recovering thawed MSCs prolongs CFT

Immobilizing MSCs in the bioreactor allows for cell recovery post-thaw, a process proposed to reduce the surface exposure of pro-coagulation factors [27]. In order to investigate the effect of recovering MSCs on clot formation, two known pro-coagulant factors-tissue factor and phosphatidylserine - were measured prior to perfusion. Flow cytometric analysis of the surface markers on both the freshly thawed and MSCs allowed to recover for 24 hours in culture, showed that cell recovery had significantly lowered the levels of phosphatidylserine and tissue factor (**Figure 3A**).

**Figure 3.**
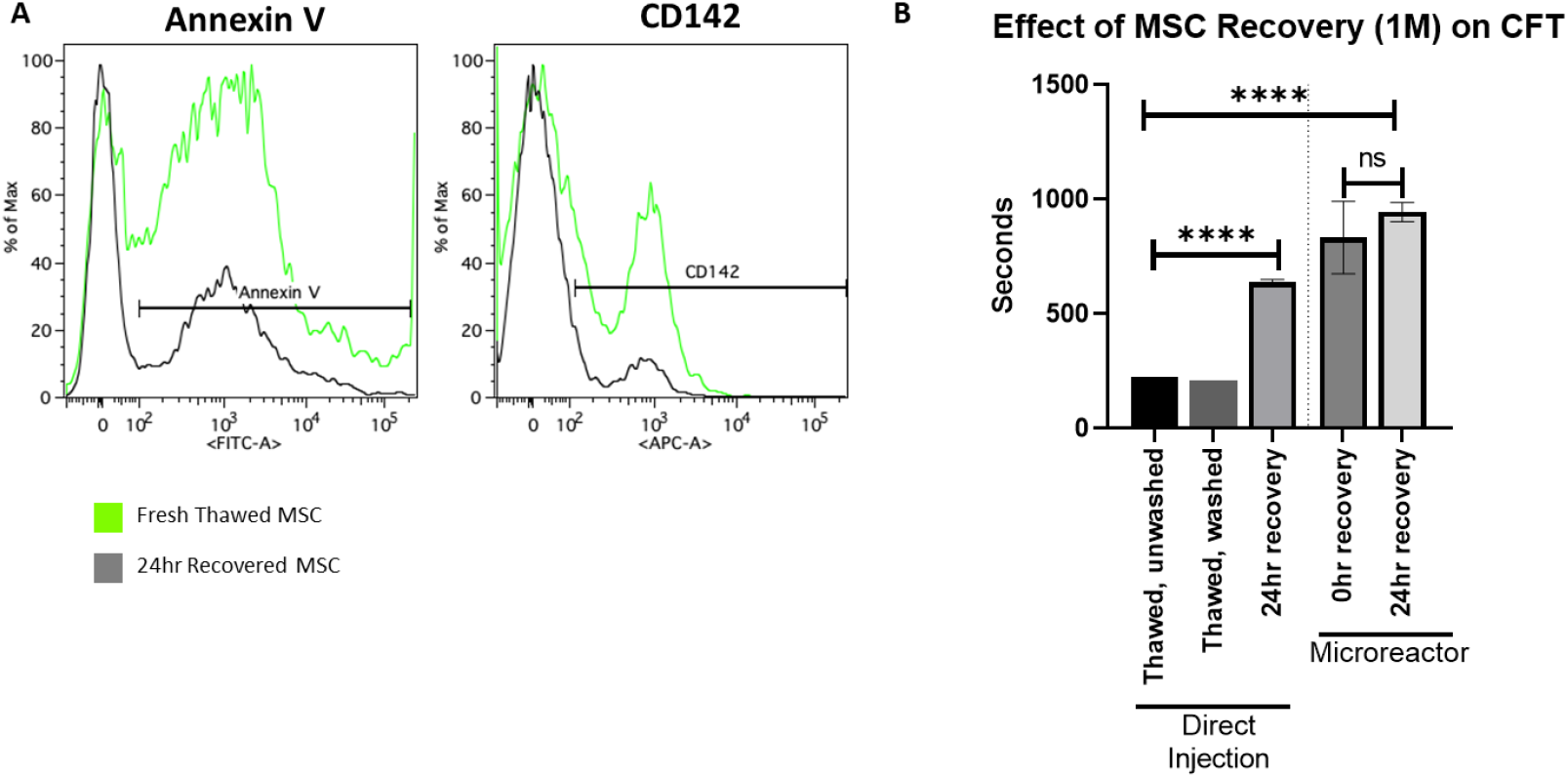
The Effect of Cell Washing and Cell Recovery on CFT. 1 x 10^6^ viable MSCs were isolated for inclusion in each respective group. Recovered cells were cultured for 24 hours and dissociated from the culture plate immediately prior to use. **(A)** Cells collected immediately after thaw and cells collected after recovery were subjected to staining and flow cytometry. Each curve represents the outcome of 3 pooled samples. **(B)** Direct injection groups were either thawed directly into plasma, washed with saline, or recovered with 24 hrs of culture at 37 °C. Microreactor groups were seeded with MSCs and either immediately used or allowed to attach for 2 hours at 37 °C followed by a 24 hour hold at room temperature prior to perfusion. Values for CFT were determined and resulting values were graphed and analyzed with an unpaired student’s t-test. N=3 runs per group. ****= p< 0.0001. Error bars represent ± standard deviation.

We next asked whether recovering MSCs post thaw had an effect on clot formation. CFTs were compared between freshly thawed cells (unwashed or washed) and cells recovered for 24 hours. Both unwashed and washed freshly thawed conditions quickly clotted at similar times during perfusion suggesting that washing to remove debris and cryopreservative did not affect clot time. However, CFT was significantly prolonged by allowing for 24 hours of recovery in culture prior to injection (**Figure 3B**).

As levels of MSC surface markers appeared to be correlated with clot initiation, we next asked whether limiting the direct interaction of MSC surface markers and plasma could reduce the rate of clot formation. Interestingly, recovery of MSCs within a microreactor did not significantly affect measured CFTs relative to fully recovered MSCs (**Figure 3B**). MSCs that were thawed, seeded into MRs, and immediately perfused, clotted at the same time as MSCs that were seeded and allowed to recover for 24 hours. These results suggest that though recovery of cells post thaw significantly reduces MSC induced clotting, housing them in an adherent state on the outside of a hollow fiber seems to further, and more significantly, reduce their clotting potential even without any recovery period.

### Removing soluble factors from the MSC reactor prolongs CFT

While seeding MSCs on the hollow fiber membrane resulted in prolonged CFT, cellular contribution was still observed as all cellular microreactor circuits clotted in a dose dependent manner (**Figure 2B**). While in the 24-hour hold period post-attachment, it is likely that MSC derived factors accumulate within the microreactor and may contribute towards clotting. To directly assay this, we integrated a saline flush of the microreactor into our protocol. After the 24-hour hold and just prior to perfusion, MRs were flushed with 3X column volume (4.5 mL) of saline. Samples from the microreactor were collected pre-and post-flush and subjected to flow cytometric analysis for measurement of tissue factor and phosphatidylserine. Pre-flush samples showed higher levels of phosphatidylserine and tissue factor in the cellular group as expected. Post-flush samples showed clear reductions in both factors, demonstrating that soluble factors can be effectively flushed out of the hollow fiber filter (**Figure 4A**). Flushing the microreactors significantly prolonged CFT in higher doses (3M), while in lower doses (1M) significance could not be reached. (**Figure 4B**). These data indicate that soluble factors (e.g. phosphotidylserine) are present in the extracapillary space of the bioreactor and can accumulate to contribute to accelerated clot formation.

**Figure 4.**
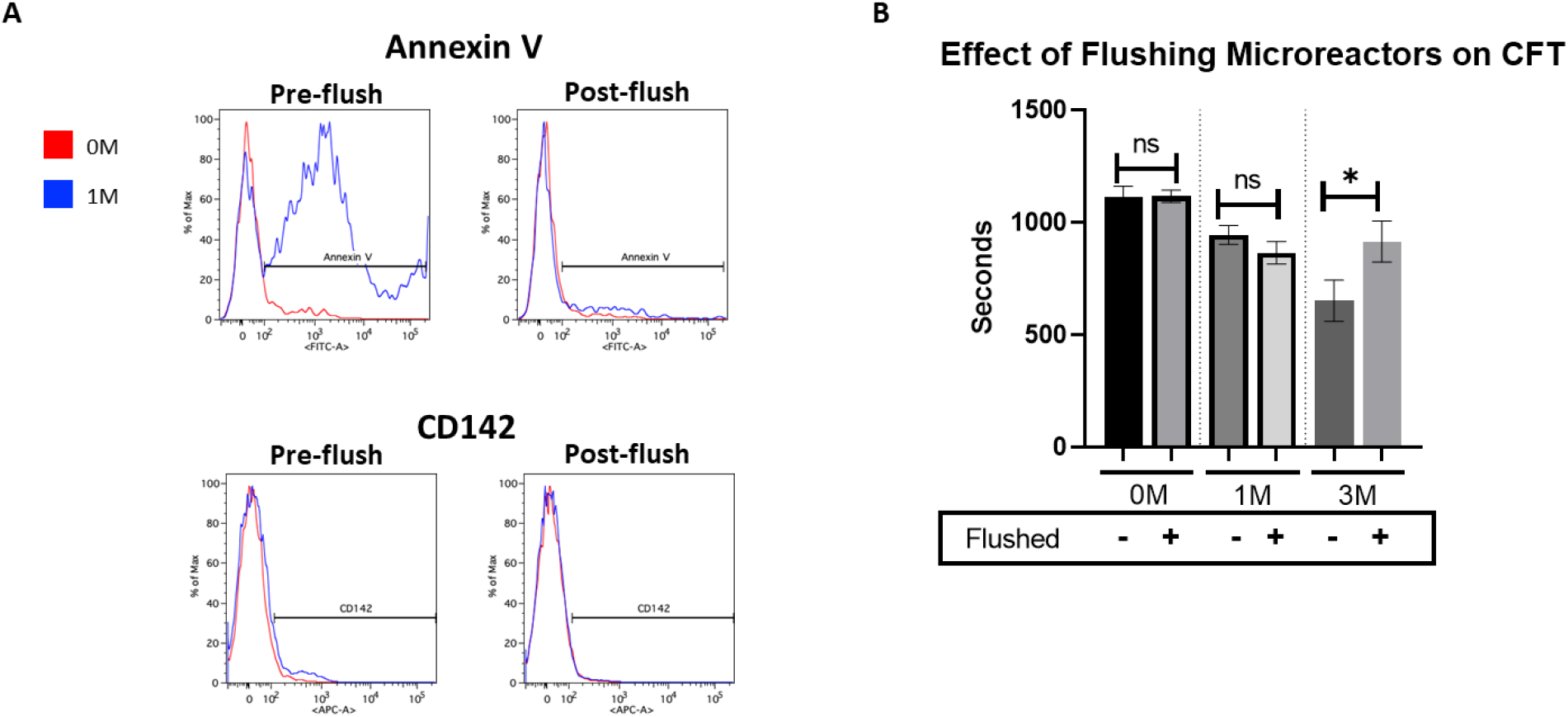
Flushing of Soluble Factors Affects CFT. Microreactors were seeded with MSCs and allowed to attach for 2 hours at 37 °C followed by a 24 hour hold at room temperature. **(A)** Samples taken from the extracapillary space of microreactors with either 0 or 1 x 10 MSCs, both pre- and post-flushing of the device, were subjected to staining and flow cytometry for known pro-coagulation markers phosphatidylserine (Annexin V) and tissue factor (CD142). Each curve represents the outcome of 3 pooled samples. **(B)** Warmed fresh frozen pooled plasma was perfused through circuits with either 0, 1 or 3 x 10^6^ viable MSCs (with and without a flushing procedure) for 5 minutes then subjected to spectrophotometric measurements. Measurements of fibrin clot formation in plasma were then made every 10 seconds over a 45-minute period. Values for CFT were determined and values were graphed and analyzed with an unpaired student’s t-test. N≥2 runs per group. *= p< 0.05. Error bars represent ± standard deviation.

### Heparin administration prevents MSC induced clotting *in vitro*

Despite significantly reducing the clotting potential of MSCs, bioreactors loaded with the cells at higher doses (3M) still induced earlier clotting when compared to acellular or low-dose (1M) cell bioreactors. Plasma spiked with heparin was used to investigate the potential efficacy of administered anticoagulation in preventing clot formation within the circuitry. Administration of 1.5 U/mL of heparin across all groups was able to completely prevent clot formation, even in the presence of positive control Innovin or 3M directly injected MSCs (**Figure 5**).

**Figure 5.**
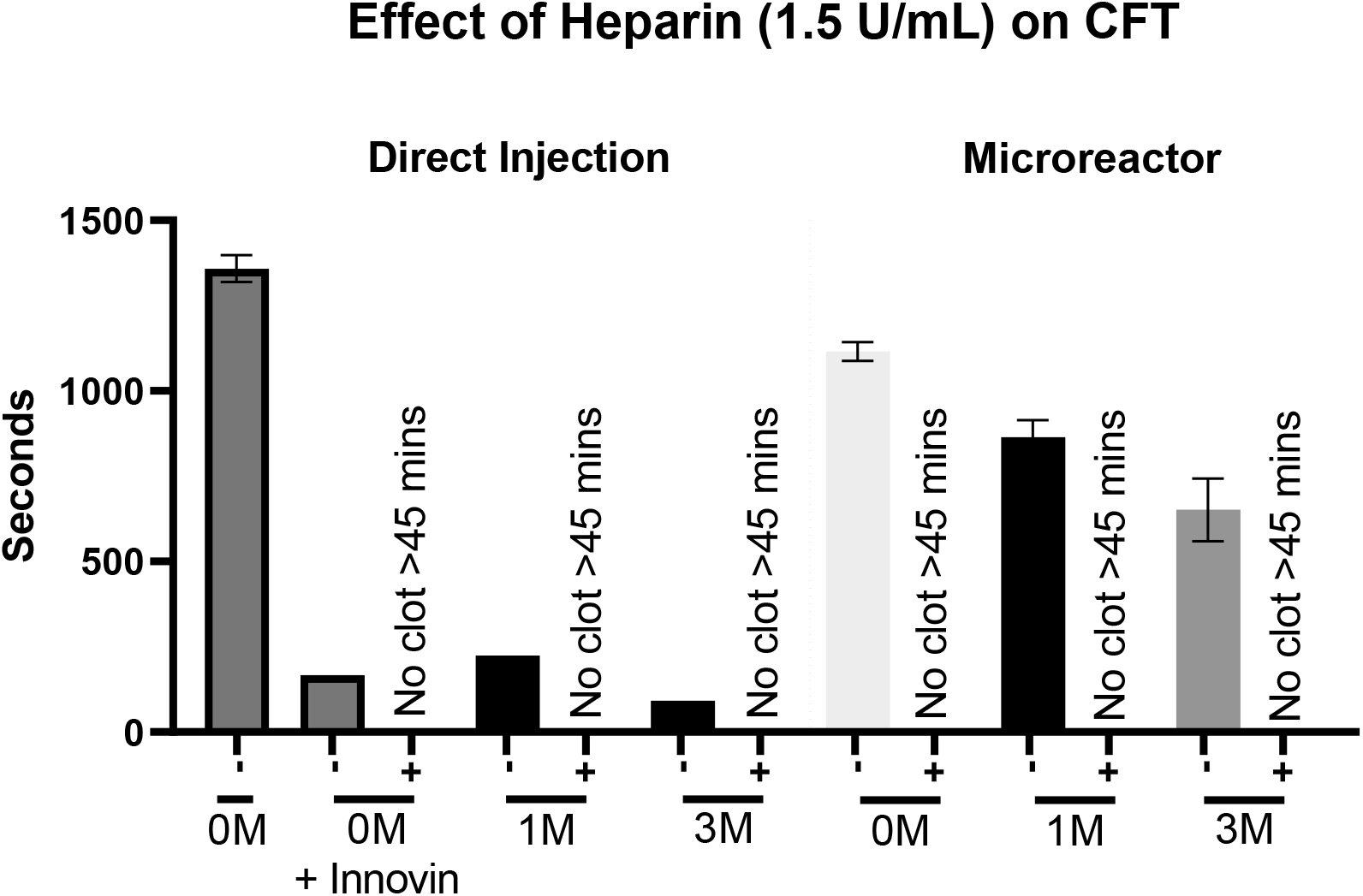
Effect of Heparin on CFT. 0, 1 x 10^6^, or 3 x 10^6^ (0M, 1M, 3M) viable MSCs were isolated for inclusion in each respective group. MSCs for direct injection groups were thawed directly into plasma then used, while microreactor groups were seeded with MSCs and allowed to attach for 2 hours at 37 °C followed by a 24 hour hold at room temperature prior to perfusion. Innovin (thromboplastin) was added to the 0M group as a positive control. After each groups’ cells were prepared, warmed FFPP was perfused through circuit for 5 minutes then subjected to spectrophotometric measurements. Measurements of fibrin clot formation in plasma were made every 10 seconds over a 45-minute period. Values for CFT were determined and resulting values were graphed and analyzed with an unpaired student’s t-test. Samples which showed no increase in absorbance through the course of the experiment were designated to have not clotted. N≥2 runs per group. *= p< 0.05. Error bars represent ± standard deviation.

### Heparin prevents MSC induced clotting *in vivo*

The bioreactor setup can be scaled up with larger filters to allow for perfusion in large animal models. Previous work in *in vivo* models showed that heparin administration could effectively reduce procoagulant activity of MSCs [35]. To reduce the number of animals used, here we compared only between bioreactor groups, no direct injection animal studies were conducted. We first tested feasibility of perfusion of the device *in vivo* in a healthy canine model. Animals were all heparinized to assure safety as extracorporeal treatments (even without cells) have intrinsic clotting potential. Dogs were grouped into cohorts based on the number of MSCs loaded into a scaled up bioreactor, with doses of 0 million, 250 million, and 750 million (n=6 dogs per group) and perfused for 24 hours. No clotting was seen in any group (data not shown).

Next, we asked the question of whether clotting in vivo would be observed under an pathological conditions, such as acute organ failure, where systemic inflammation may perturb the coagulation pathways. For this purpose a porcine animal model of acute myocardial infarction (AMI) was used (**Figure 6 A**). AMI was induced, animals were re-perfused/stabilized for 1 hour and then connected to the bioreactor perfusion circuit for 12 hours. All animals were perfused without events for 12 hours, with each group showing cardiac injury biomarker induction (**Figure 6 B**) and similar infarct size (**Figure 6 C**). Heparin was administered throughout the perfusion process to maintain a minimum activated clotting time (ACT) of at least 300 seconds (as mandated by IACUC), with neither group requiring significantly more heparin than the other (**Figure 6 D, E**). These data support the use of MSC bioreactors without additional heparin requirements beyond what is used in acellular extracorporeal treatments.

**Figure 6.**
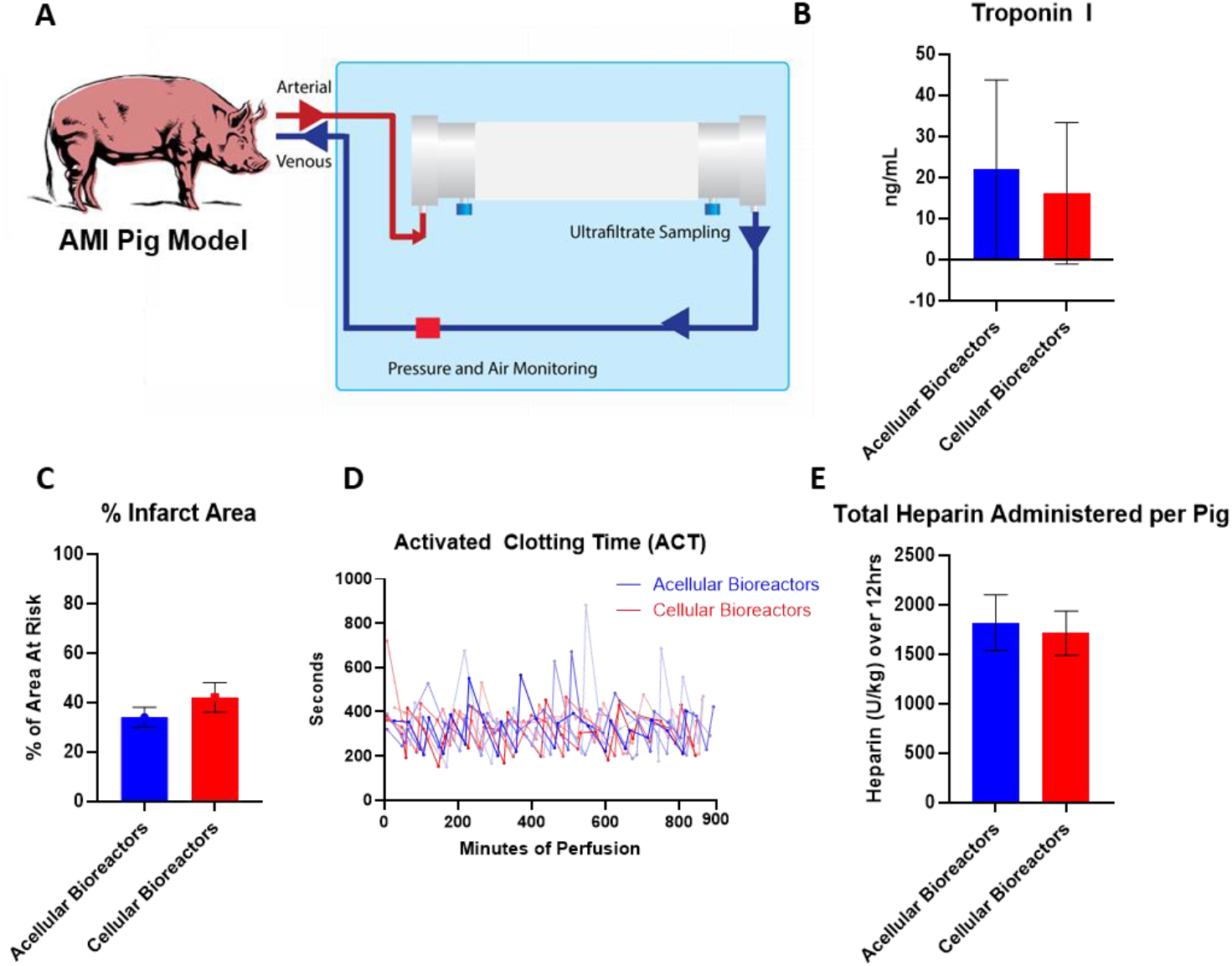
Pig AMI Model Perfusion. **(A)** Pigs were sedated, occluded of their left anterior descending coronary artery, re-perfused for 1 hour, and administered the designated bioreactor (acellular or cellular) for 12 hours of perfusion. **(B,C)** Comparative measurements of induced stress were done through serum sampling of Troponin I levels at the 12 hour mark, as well as morphological analysis of % infarct area at sacrifice. **(D,E)** Heparin was administered over the course of 12 hours to maintain an ACT of approximately 300 seconds. N=4 pigs per group. Error bars represent ± standard deviation.

## Discussion

In the absence of clear clinical benefit of early allogeneic MSC human trials to meet their therapeutic endpoints, there has been a major focus in recent years to improve the reliability and consistency of the therapeutic cells delivered [40]. Improvements in manufacturing processes, more stringent release testing and biobanking has provided a reproducibility to the cell production that has contributed to a clinically approved therapy [8, 28, 41–43]. However, many cellular concerns still exist, including handling at point of care, thawing, route of delivery, hemocompatibility, and dosing. Our studies here focused on comparing the potential risks of one of those concerns, MSC induced coagulation, between direct infusion and a modified ex vivo, systemic approach.

Most commonly, allogeneic MSCs are delivered for therapeutic effect through systemic administration (intravenous or intra-arterial) accounting for about half of all published studies [28]. Systemic introduction has been described as the least invasive, most reproducible, and provides the MSCs the most direct access to modulate systemic inflammation [44]. However, this route of administration may increase risks to certain hypercoagulable patients given that MSCs are known to express coagulation factors both on their cell surface and on the exosomes and vesicles they secrete, namely tissue factor and phosphatidylserine [16, 41, 45]. Further, systemically introduced MSCs can rapidly get trapped in the lungs or be cleared, reducing their potential efficacy [40]. Because of these concerns, where possible and concordant with the mechanism of action (MoA), alternatives to systemic administration are increasingly utilized, including intramuscular infusions, topical, direct tissue injections, and intracoronary delivery. While useful for localized applications including tissue regeneration, these routes of delivery are not used to treat systemic applications such as GvHD, and present limitations of their own in terms of feasibility, reproducibility, and efficacy [46, 47].

Here, in concert with the previously mentioned improvements with cellular production, we assayed the value of incorporating an experimental setup which confines MSCs behind the membrane of a hollow-fiber filter. Given that much of the MSCs’ ability to induce clot formation arises from its cell surface markers and secreted vesicles, we considered that the confinement of the cells and their procoagulant expressing surface markers in one extraluminal location may reduce the rate at which clot formation occurs. We used an existing bioreactor platform known to retain MSC immunomodulatory capacity in combination with modifications to an existing clot formation assay to assess cellular effect on CFT in this immobilized state relative to direct injection [32, 34, 35]. Through this platform we perfused citrated, platelet poor fresh-frozen pooled plasma. This plasma contains many essential factors for clot initiation, including prothrombin which can be activated with the addition of calcium via CaCl_2_ to form firm clots over time.

Immediately after CaCl_2_ addition, this coagulable (but still liquid) plasma is perfused through the hollow-fiber filter platform. As expected, direct injection of MSCs into the coagulable plasma perfusion circuit led to rapid clot formation in a dose-dependent manner. Interestingly, perfusion of coagulable plasma through bioreactors seeded with MSCs resulted in clotting at rates significantly slower than their comparable direct injection groups, suggesting that free, circulating MSCs increase thrombosis risk more than bioreactor immobilized MSCs. Like the direct injection group, the MSC dose seeded in the bioreactor was predictive of CFT with higher doses inducing quicker clots, likely through the increased production of pro-coagulable MSC factors. It is also important to note that the presence of a filter (acellular microreactor) in the circuit induced clotting faster than a circuit without a microreactor supporting the idea that high surface area biomaterials increase factor adsorption and may contribute to expedited clotting [48]. Alternative approaches such as heparin coating the hollow fiber filters prior to MSC seeding may reduce surface adsorption and adhesion, lowering this inherent clot induction potential [49–51].

Historically, failed MSCs trials have been in part attributed to poor cell processing, including delivery of dead and/or coagulable cells [40, 43, 52, 53]. In this study, washing of cells post thaw did not affect CFT significantly. However, recovery of cells for 24 hours post thaw did reduce clot formation potential. This was shown to correlate with surface marker expression of tissue factor and phosphatidylserine. Both decreased following recovery, correlating with a slower clot formation time relative to MSCs directly injected into the circuit post thaw. Recovery culture of cryopreserved MSCs within a microreactor did not have a significant impact on CFT relative to freshly thawed cells within microreactors, suggesting that the MSC confinement by the hollow fiber membrane may actually be playing a role in prolonging CFT. While MSCs located behind the membrane are still able to exchange their immunomodulatory secreted factors with perfusing solutions, they may be sharing less of their cell surface area and may aid in confining their cellular debris to the extracapillary space of the microreactor. Future studies with more restrictive filter sizes may even further limit MSC exposure and further prolong CFT.

Having shown that cell presence shortens clot formation time we sought to use the platform to mitigate that effect as much as possible. Since our microreactor is composed of hollow fibers, exchange does happen through the 0.2 μm pores on the fibers in the microreactor. During hold, MSCs continue to produce materials and some of this accumulated material could be contributing to clot formation. To assess this, we developed a flush protocol and measured steep drops in known coagulation markers. Consequently, flushing resulted in slower CFTs at higher MSC doses. Future studies will assay whether flushing also affects immunomodulatory potential relative to unflushed reactors [32].

Despite the microreactors measured effect of prolonging CFT, it did not completely abrogate the cellular contribution to shortened CFT. In clinical setting anticoagulation protocols will likely be integrated to ensure designated perfusion times are met. Our *in vitro* experiments and *in vivo* canine studies showed that heparin administration could effectively prevent any cellular induced clot formation during perfusion. However, many of the patients suffering from systemic inflammation, including those with COVID-19, present with hypercoagulable plasma that will require anticoagulation prior to MSC administration. Our pig model represented a more physiologically relevant condition in which inflamed animals were perfused with a device scaled for human use. Under these acute injury conditions, no clotting was observed in animals perfused with devices loaded with 750M MSCs and for 12 hours. The lack of additional heparin requirement suggests that patients set to undergo MSC-bioreactor perfusion may not need more heparin than a sham control undergoing the same procedure. Future studies comparing direct infusion of MSCs to bioreactor housed MSCs would be useful to evaluate both for coagulation and efficacy responses.

Given the slower CFT in the microreactor groups relative to the direct injection groups, it is possible that a lower dose of heparin could be administered to the microreactor groups. Future dose testing will be required to verify this. Such a finding would be clinically relevant, as reduction in the amount of heparin required to be delivered to critically ill patients may help prevent unintended health consequences. Further, patients which are medically restricted from systemic heparin administration for risk of internal bleeding could potentially be anticoagulated regionally with citrate. Citrate could be introduced and equilibrated within the MSC bioreactor circuit, allowing MSC factors to be released but without exposing patients to the anticoagulant [54].

In the unfortunate circumstances of the COVID-19 pandemic, interest in MSC based therapies has increased markedly. Case reports, first from China and then worldwide, showed promising improvements in patient health following intravenous MSC infusion, even in severely ill patients [10, 55, 56]. While larger studies are now needed to more completely support these findings, it is clear that intravenous infusion of MSCs for systemic inflammatory conditions such as COVID-19 infection or GvHD continues to have therapeutic potential. Remestemcel-L, an ex-vivo culture-expanded adult human MSC suspension for intravenous infusion, which has received positive recommendations from the FDA for steroid-refractory acute graft-versus-host disease in pediatric patients based on an open-label study compared to historical controls [57]. The novel delivery approach described here could potentially reduce risk of clot formation from IV administered MSCs, making treatment potentially safer and more controlled than direct infusion.

## Conclusion

We conclude that immobilization of MSCs in a hollow fiber filter contributes to a reduced clot initiation potential relative to directly injected MSCs. Further removal of cellular byproducts through saline flushing of the bioreactor further reduces the MSC based clot formation potential. Additional heparin does not appear to be required to maintain a designated ACT value relative to acellular perfusion circuits. Taken together, combined integration of these approaches may make MSC therapies which require systemic MSC exposure at less risk for coagulation-related events for a larger population, including the hypercoagulable.

## Disclosures of Potential Conflicts of Interest

The opinions or assertions contained herein are the private views of the author and are not to be construed as official or as reflecting the views of the Department of the Army or the Department of Defense.

JB and AC are United States government employees with no financial disclosures relevant to this publication.

BOR, SN, AT, RNB are employees and equity shareholders of Sentien Biotechnologies. BP is an equity shareholder and inventor of the technology with licensed patents to Sentien for commercialization.

## Funding

This research was conducted with private funding.

## Data Availability Statement

The data that support the findings of this study are available from the corresponding author upon reasonable request.

## Notes

### Summary of Updates

The title was revised slightly for clarity.

