## Supplemental Figures for "Mesenchymal Stromal Cell Delivery via an Ex Vivo Bioreactor Preclinical Test System Attenuates Clot Formation for Intravascular Application"

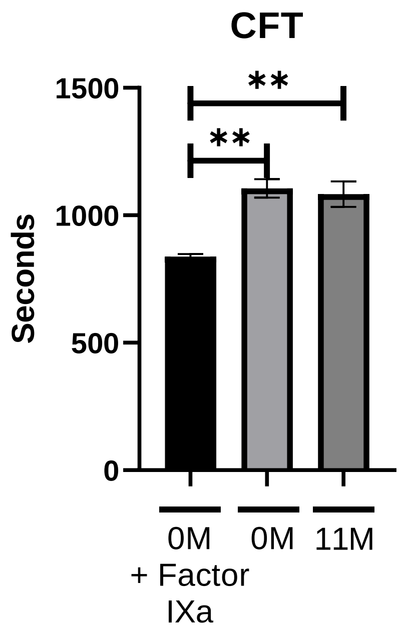


**Figure S1. Single Donor Plasma Insensitivity.** Microcreactors were seeded with either 0 or 11 x $10^6$ viable MSCs and allowed to attach for 2 hours at 37 °C, followed by a 24 hour hold at room temperature prior to perfusion. Factor IXa was added to the 0M group as a positive control of clot formation. After each groups' cells were prepared, warmed FFPP was perfused through circuit for 5 minutes then subjected to spectrophotometric measurements. (A) Measurements of fibrin clot formation in plasma were then made every 10 seconds over a 45-minute period. Values for CFT were determined and resulting values were graphed and analyzed with an unpaired student's t test. N≥2 runs per group. **= $p< 0.005$. Error bars represent ± standard deviation.

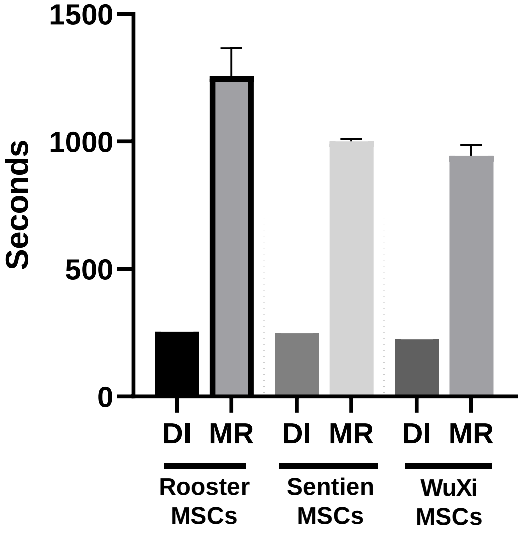


**Figure S2. MSC donor testing.** Microcreactors were seeded with 1 x $10^6$ viable MSCs each from distinct donors and allowed to attach for 2 hours at 37 °C, followed by a 24 hour hold at room temperature prior to perfusion. After each groups' cells were prepared warmed FFPP was perfused through circuit for 5 minutes then subjected to spectrophotometric measurements, if not already clotted. (**A**) Measurements of fibrin clot formation in plasma were then made every 10 seconds over a 45-minute period. Error bars represent ± standard deviation.
